## Supporting Information for "Influenza A virus resistance to 4’-fluorouridine coincides with viral attenuation *in vitro* and *in vivo*"

**S1 Table.** Whole genome sequencing of adapted virus populations after passage 9 (P9) or 10 (P10) as indicated. Shown are coding mutations that were absent in lineages passaged in the presence of vehicle (DMSO) volume equivalents. Allele frequency in parentheses; frequency cut-off 5%; read depth cut-off 50. Bold underscore denotes mutations in RdRP subunits with >50% relative allele frequency that were rebuilt for resistance testing.

| adaptation lineage | M | HA | NA | NP | NS | PA | PB1 | PB2 |
|---|---|---|---|---|---|---|---|---|
| #1<br>(after P9) | | | V53F<br>(32.76%) | D72A<br>(5.98%)<br>S430N<br>(46.65%) | I54T<br>(99.43%) | M374V<br>(46.46%) | **V285I<br>(97.63%)** | |
| #2<br>(after P9) | | | G11S<br>(5.99%)<br>N63D<br>(29.87%) | D72A<br>(5.71%) | | | **T46A<br>(99.88%)** | N71D<br>(10.2%)<br>**E180K<br>(57.39%)**<br>**E191K<br>(59.21%)**<br>M483K<br>(31.02%)<br>V667I<br>(22.69%) |
| #3<br>(after P9) | | | | D72A<br>(5.61%) | A89E<br>(7.31%) | S291G<br>(5.28%) | D76N<br>(36.1%)<br>**M290V<br>(50.67%)**<br>I392V<br>(12.61%) | **K189R<br>(100%)** |
| #4<br>(after P10) | | E85G<br>(98.34%) | Y66C<br>(12.48%) | V186I<br>(15.49%) | I54T<br>(86.94%) | E26G<br>(17.16%)<br>**S395N<br>(72.96%)** | | **Y488C<br>(99.1%)**<br>**T491M<br>(69.52%)** |
| #5<br>(after P9) | | A152S<br>(80%) | | D72A<br>(6.01%)<br>S84N<br>(21.07%) | M104I<br>(7%) | **N222S<br>(71.63%)**<br>D386N<br>(18.68%) | **V285I<br>(99.92%)** | T16I<br>(9.67%)<br>S92A<br>(24.24%)<br>A271T<br>(5.37%)<br>D408G<br>(5.94%)<br>V414I<br>(32.77%)<br>D455Y<br>(32.45%) |
| #6<br>(after P10) | | | S31P<br>(99.94%)<br>P126L<br>(7.49%) | | | **M579I<br>(81.87%)** | **M339I<br>(99.97%)** | **Y488C<br>(98.5%)**<br>V731I<br>(32.77%) |

**S2 Table:** Dose response assays of recCA09 with rebuilt resistance mutations against 4'-FIU ($EC_{99}$ with 95% CI and fold-change $EC_{99}$ relative to parental recCA09 are shown).

| adaptation lineage | mutation | $EC_{99}$ and 95% CI | fold-change |
|---|---|---|---|
| WT | | 0.53 µM (0.181 - 1.703) | N/A |
| #1 | PB1 (V285I) | 0.96 µM (0.375 - 4.691) | 2× |
| #2 | PB1 (T46A) + PB2 (E180K, E191K) | 2.98 µM (0.731 - 19.78) | 6× |
| #3 | PB1 (M290V) + PB2 (K189R) | 9.19 µM (1.91 - 71.74) | 17× |
| #4 | PA (S395N) + PB2 (Y488C, T491M) | 7.75 µM (1.784 - 42.82) | 15× |
| #5 | PA (N222S) + PB1 (V285I) | 13.15 µM (4.649 - 44.87) | 25× |
| #6 | PA (M579I) + PB1 (M339I) + PB2 (Y488C) | 1.27 µM (0.2 - 20.28) | 2× |
| favipiravir resistant | PB1 (K229R) + PA (P653L) | 0.52 µM ( - 4.97) | 1× |

**S3 Table:** Dose response assays of recCA09 with rebuilt resistance mutations against favipiravir/T-705 ($EC_{99}$ with 95% confidence CI and fold-change $EC_{99}$ relative to parental recCA09 are shown).

| adaptation lineage | mutation | $EC_{99}$ and 95% CI | fold-change |
|---|---|---|---|
| WT | | 2.96 µM ( - 15.73) | N/A |
| #1 | PB1 (V285I) | 7.81 µM (2.21 - 34.78) | 3× |
| #2 | PB1 (T46A) + PB2 (E180K, E191K) | 5.52 µM (0.373 - 284.5) | 2× |
| #3 | PB1 (M290V) + PB2 (K189R) | 6.32 µM (0.236 - 660.6) | 2× |
| #4 | PA (S395N) + PB2 (Y488C, T491M) | 35.45 µM ( - 1521) | 12× |
| #5 | PA (N222S) + PB1 (V285I) | 11.09 µM (1.769 - 86.47) | 4× |
| #6 | PA (M579I) + PB1 (M339I) + PB2 (Y488C) | 9.23 µM (1.293 - 112.4) | 3× |

**S4 Table:** Dose response assays of recCA09 with rebuilt resistance mutations against NHC (parent compound of the prodrug molnupiravir; $EC_{99}$ with 95% confidence CI and fold-change $EC_{99}$ relative to parental recCA09 are shown).

| adaptation lineage | mutation | $EC_{99}$ and 95% CI | fold-change |
|---|---|---|---|
| WT | | 3.38 µM ( - 23.28) | N/A |
| #1 | PB1 (V285I) | 0.32 µM ( - 4.570) | 0.1× |
| #2 | PB1 (T46A) + PB2 (E180K, E191K) | 1.01 µM (0.206 - 11.52) | 0.3× |
| #3 | PB1 (M290V) + PB2 (K189R) | 2.29 µM ( - 983.6) | 0.7× |
| #4 | PA (S395N) + PB2 (Y488C, T491M) | 0.9 µM ( - 120.6) | 0.3× |
| #5 | PA (N222S) + PB1 (V285I) | 0.31 µM ( - 1.849) | 0.1× |

| | | | |
|---|---|---|---|
| #6 | PA (M579I) + PB1 (M339I) + PB2 (Y488C) | 2.61 μM (0.862 - 10.34) | 0.8× |

**S5 Table:** Peak virus titers and maximum growth rates of recCA09 with rebuilt resistance mutations.

| adaptation lineage | mutation | max growth rate [$\mu_{max/h}$] | max titer [$TCID_{50}$/ml] |
|---|---|---|---|
| WT | | 0.22 | $1.05\times10^{6}$ |
| #1 | V285I | 0.19 | $6.49\times10^{4}$ |
| #2 | T46A +E191K+E180K | 0.21 | $3.79\times10^{5}$ |
| #3 | M290V+K189R | 0.25 | $3.1\times10^{5}$ |
| #4 | S395N+Y488C+T491M | 0.24 | $8\times10^{5}$ |
| #5 | N222S+V285I | 0.19 | $6.49\times10^{4}$ |
| #6 | M579I+M339I+Y488C | 0.22 | $1.93\times10^{5}$ |

**S6 Table:** Representation of candidate resistance mutations in complete and partial IAV sequences available in the NIH NCBI Virus sequence database (limit: 0.001%). Polymorphisms matching resistance mutations are shown in bold.

**Resistance mutations adaptation: N222S; S395N; M579I**

| PA | complete sequences (1147) | partial sequences (96048) |
|---|---|---|
| N222S | N 99.04%<br>**S 0.7%**<br>H 0.17%<br>K 0.09% | N 98.549%<br>**S 0.094%**<br>H 0.036%<br>K 0.033%<br>D 0.012%<br>Y 0.011%<br>G 0.006%<br>E 0.001%<br>T 0.001% |
| S395N | S 98.61%<br>C 1.31%<br>**N 0.09%** | S 98.814%<br>**N 0.203%**<br>G 0.119%<br>C 0.069%<br>T 0.049%<br>H 0.009%<br>P 0.006%<br>I 0.002%<br>R 0.001% |
| M579I | M 99.74%<br>**I 0.17%**<br>V 0.09% | M 98.862%<br>**I 0.167%**<br>V 0.04%<br>T 0.006%<br>L 0.005% |

**Resistance mutations adaptation: T46A, V285I, M290V, M339I**

| PB1 | complete sequences (1139) | Partial sequences (94791)p |
|---|---|---|
| T46A | T 99.65%<br>Q 0.18% | T 98.926%<br>**A 0.001%** |

| | | |
|---|---|---|
| | R 0.18% | G 0.002%<br>R 0.002%<br>K 0.001% |
| V285I | V 99.39%<br>Y 0.44%<br>R 0.18% | V 98.557%<br>A 0.002%<br>L 0.001% |
| M290V | M 99.39%<br>Q 0.44%<br>N 0.18% | M 98.521%<br>I 0.012%<br>T 0.005%<br>**V 0.003%**<br>E 0.001%<br>L 0.001% |
| M339I | **I 71.47%**<br>M 27.22%<br>V 0.7%<br>L 0.44%<br>P 0.18% | **I 71.529%**<br>M 25.951%<br>V 1.019%<br>L 0.011%<br>T 0.008%<br>F 0.002%<br>S 0.002%<br>K 0.001% |

**Resistance mutations adaptation: E180K, K189R, E191K, Y488C, T491M**

| PB2 | complete sequences (1182) | partial sequences (96301) |
|---|---|---|
| E180K | E 99.92%<br>A 0.08% | E 98.346%<br>D 0.077%<br>G 0.015%<br>**K 0.007%**<br>A 0.001% |
| K189R | K 99.92%<br>**R 0.08%** | K 98.476%<br>**R 0.01%**<br>I 0.002%<br>N 0.002%<br>S 0.002%<br>M 0.001%<br>Q 0.001%<br>T 0.001% |
| E191K | E 95.44%<br>**K 4.48%**<br>G 0.08% | E 96.652%<br>**K 1.137%**<br>D 0.573%<br>G 0.079%<br>A 0.017%<br>R 0.012%<br>N 0.011%<br>Q 0.006%<br>V 0.004%<br>L 0.002%<br>M 0.001%<br>S 0.001%<br>T 0.001% |
| Y488C | Y 99.92%<br>F 0.08% | Y 98.665%<br>H 0.036%<br>F 0.012%<br>**C 0.005%**<br>D 0.001% |
| T491M | T 99.66%<br>A 0.34% | T 95.598%<br>A 2.039%<br>I 0.058%<br>N 0.008%<br>S 0.005%<br>**M 0.004%**<br>E 0.001% |

| | | |
|---|---|---|
| | | P 0.001%<br>R 0.001%<br>V 0.001% |

**S7 Table:** Location and predicted effect of individual resistance mutations.

| adaptation lineage | PA | PB1 | PB2 | rationale/location |
|---|---|---|---|---|
| #1 | | V285I | | Mutation adds bulk and pushes on the base of motif C which may alter the geometry of the polymerase active site. |
| #5 | N222S | V285I | | |
| | | V285I | | Mutation adds bulk and pushes on the base of motif C which may alter the geometry of the polymerase active site. |
| | N222S | | | Removes bulk from a central PA/PB1 interface that allows for changes in the interior channels of the polymerase. |
| #2 | | T46A | E180K<br>+<br>E191K | |
| | | T46A | | Removes bulk and sits adjacent to motifs A and B. Might alter the position of the GDN site or alter the arrangement of RNA within the RNA cavity. |
| | | | E180K<br>+<br>E191K | Alters structural mobility of PB2, allowing for altered loading of cap RNA into the central cavity. |
| #3 | | M290V | K189R | |
| | | M290V | | Mutation reduces bulk and might allow for PB2 to reach further into core of RNA pol during cap addition, thus altering the interior environment of the RNA cavity. |
| | | | K189R | Alters structural mobility of PB2, allowing for altered loading of cap RNA into the central cavity. |
| #4 | S395N | | Y488C<br>+<br>T491M | |
| | S395N | | | Sits by the RNA binding site for template RNA loop. No direct effect on resistance, but may alter the loading of template RNA, working with the Y488C mutation to change geometry of RNA cavity. |
| | | | Y488C | Located on PB2 and is positioned close to V285I when the cap binding domain of PB2 is brought towards the RNA cavity for cap priming. Cooperation with T491M to change the positioning of PB2 bound cap intermediate to allow for altered NTP selectivity through changes in RNA conformation within the interior cavity. |

| | | | | |
|---|---|---|---|---|
| | | | T491M | Gains bulk and may work in cooperation with Y488C to change the positioning of PB2 bound cap intermediate to allow for altered NTP selectivity through changes in RNA conformation within the interior cavity. |
| #6 | M579I | M339I | Y488C | |
| | M579I | | | Removes bulk from a central intersection of PA, PB1, and PB2. This may lead to altered structure of the interior cavity. |
| | | M339I | | Near Motif F. Mutation may alter positioning of motif F, potentially allowing for altered selectivity. Sits on the opposite side of the polymerase compared to Y488C and might work in tangent to alter RNA channel and catalytic site geometry. |
| | | | Y488C | Located on PB2 and is positioned close to V285I when the cap binding domain of PB2 is brought towards the RNA cavity for cap priming. Might work with M339I mutation. |

**S8 Table:** Recovery attempts of recCA09 with engineered combinations of independently emerged resistance mutations (genetic background: recCA09-nanoLuc).

| virus | attempt 1 (11/15/22) | attempt 2 (11/27/22) | attempt 3 (12/10/22) | attempt 4 (12/20/22) |
|---|---|---|---|---|
| S395N + Y488C | no recovery | no recovery | no recovery | no recovery |
| T491M | no recovery | no recovery | no recovery | no recovery |
| S395N + T491M | no recovery | no recovery | no recovery | no recovery |
| V285I + K189R | no recovery | no recovery | no recovery | no recovery |
| N222S + M290V | no recovery | no recovery | no recovery | no recovery |
| V285I + Y488C | no recovery | no recovery | no recovery | no recovery |
| S395N + V285I + Y488C + T491M | no recovery | no recovery | no recovery | no recovery |
| WT | recovery | recovery | recovery | Recovery |

### Supporting information captions

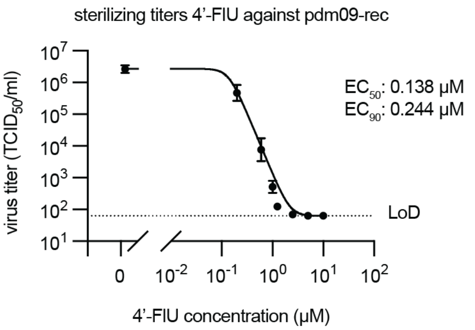


**S1 Figure:** Sterilizing dose-range finding with 4'-FIU. Virus yield reduction assay with recCA09. Symbols represent geometric mean □ geometric SD; line shows 4-parameter variable slope regression model. $EC_{50}$ and $EC_{90}$ values are given; LoD, Limit of Detection; n=3.

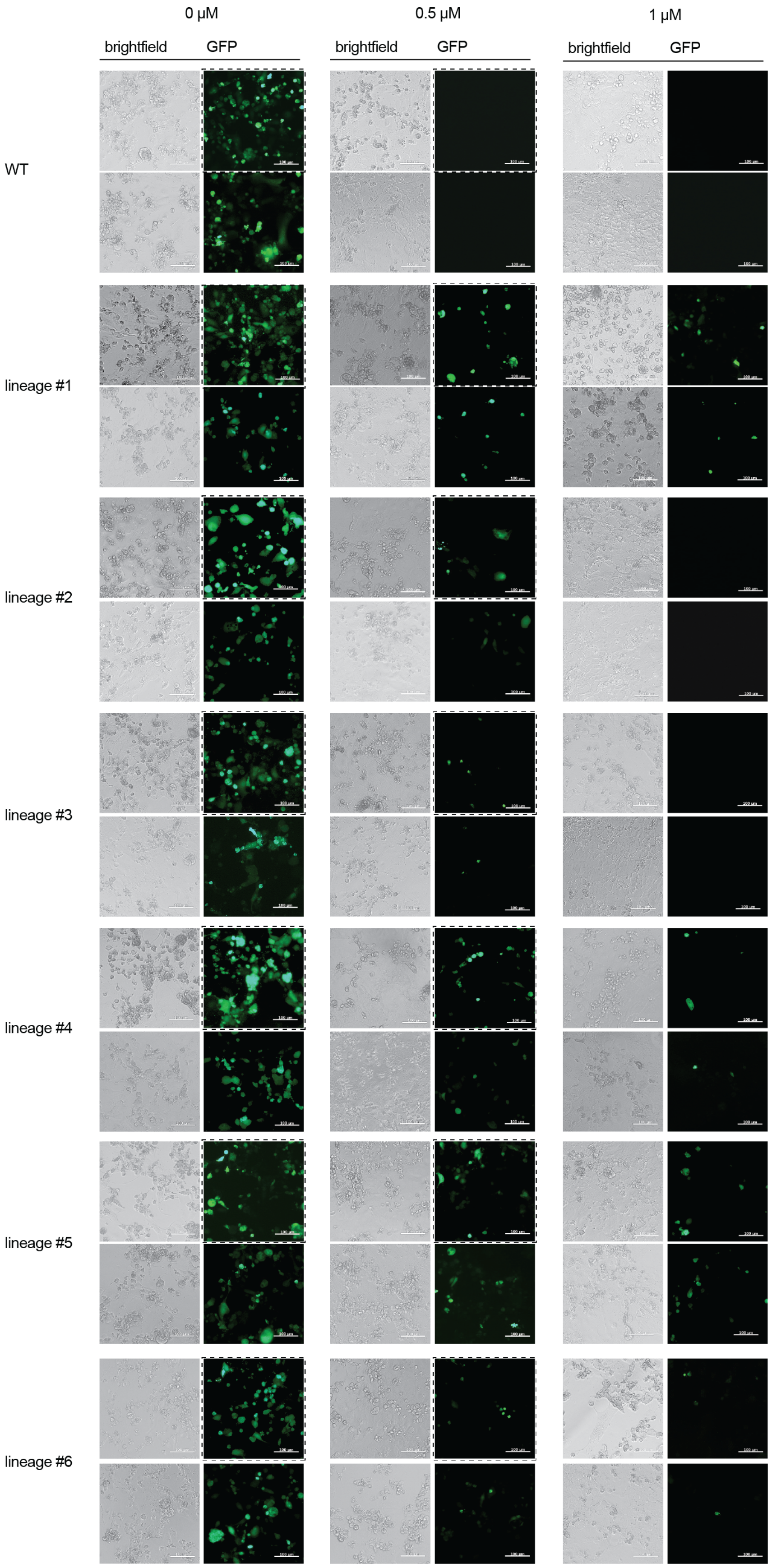
0 µM
0.5 µM
1 µM
brightfield
GFP
brightfield
GFP
brightfield
GFP
WT
lineage #1
lineage #2
lineage #3
lineage #4
lineage #5
lineage #6

44

**S2 Figure:** Repeats of microphotographs shown in (Fig. 1e). Duplicates of phase-contrast images and corresponding fluorescent images are shown for each adaptation lineage; dashed boxes denote images presented in Fig 1e; scale bar, 100 µm.

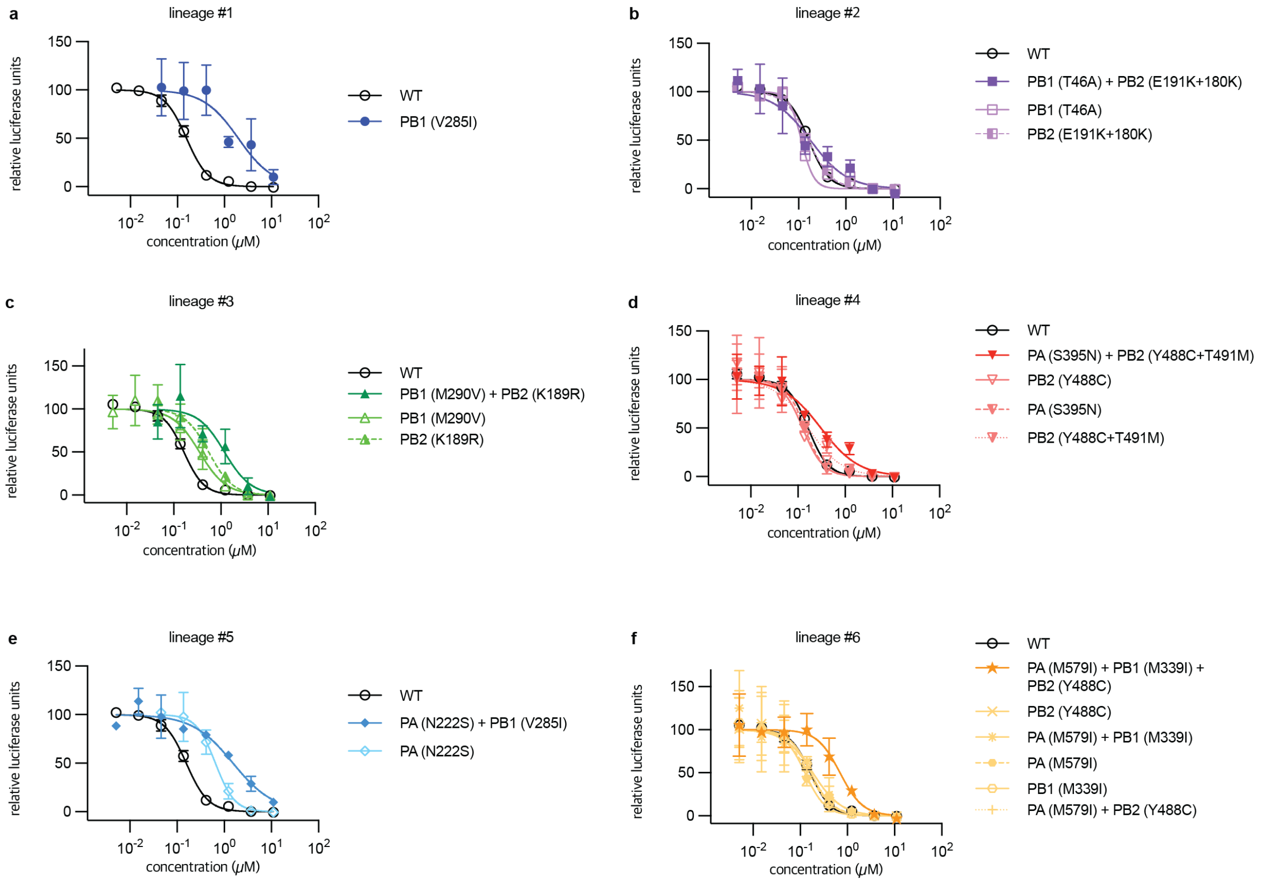


**S3 Figure:** Dose-response assays of recCA09-nanoLuc with resistance mutations rebuilt in combination and individually. **a-f)** Assessment of recCA09 with mutations found in lineages 1 (a), 2 (b), 3 (c), 4 (d), 5 (e), and 6 (f). Symbols show means □ SD, lines show 4-parameter variable slope regression models. Data were normalized for samples receiving vehicle (DMSO) volume equivalents, parental recCA09 is shown in each graph; n=3.

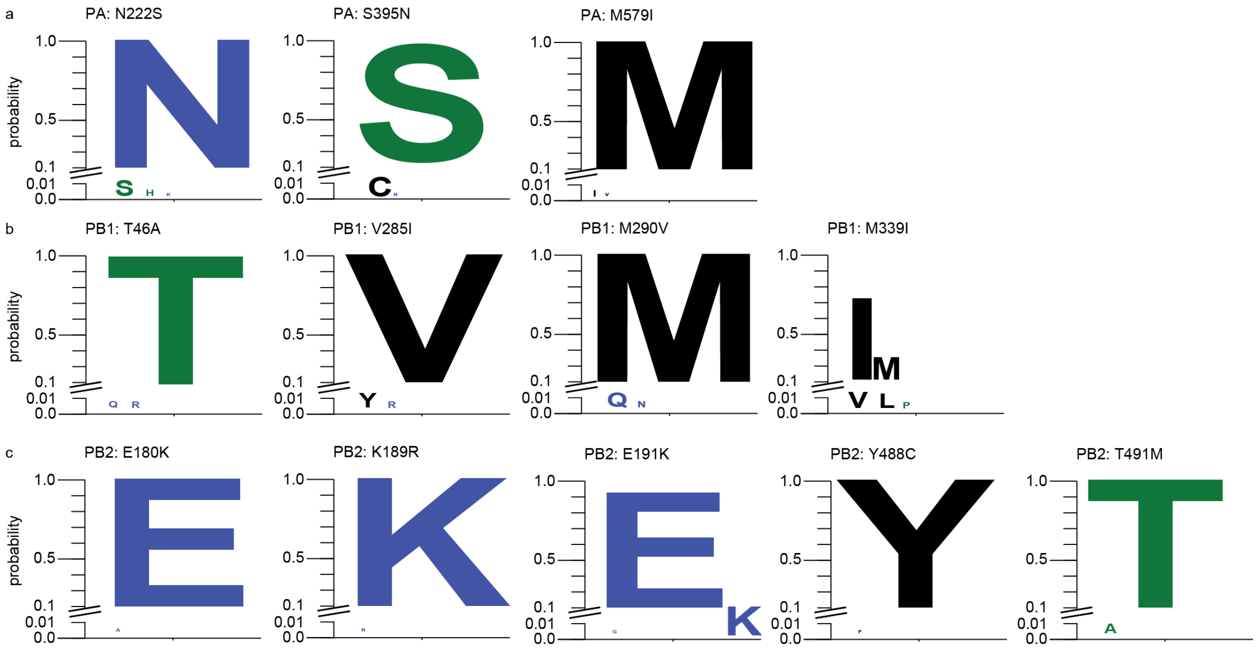


**S4 Figure:** Representation of candidate resistance mutations in complete IAV sequences available in the NIH NCBI Virus sequence database. **a-c)** Shown are relative frequency of polymorphism at 4'-FlU resistance sites in PA (a), PB1 (b), and PB2 (c). Blue, predominant hydrophilic site chains; green, predominant neutral site chains; black, predominant hydrophobic site chains. Relative size proportional to probability of presence in the database.

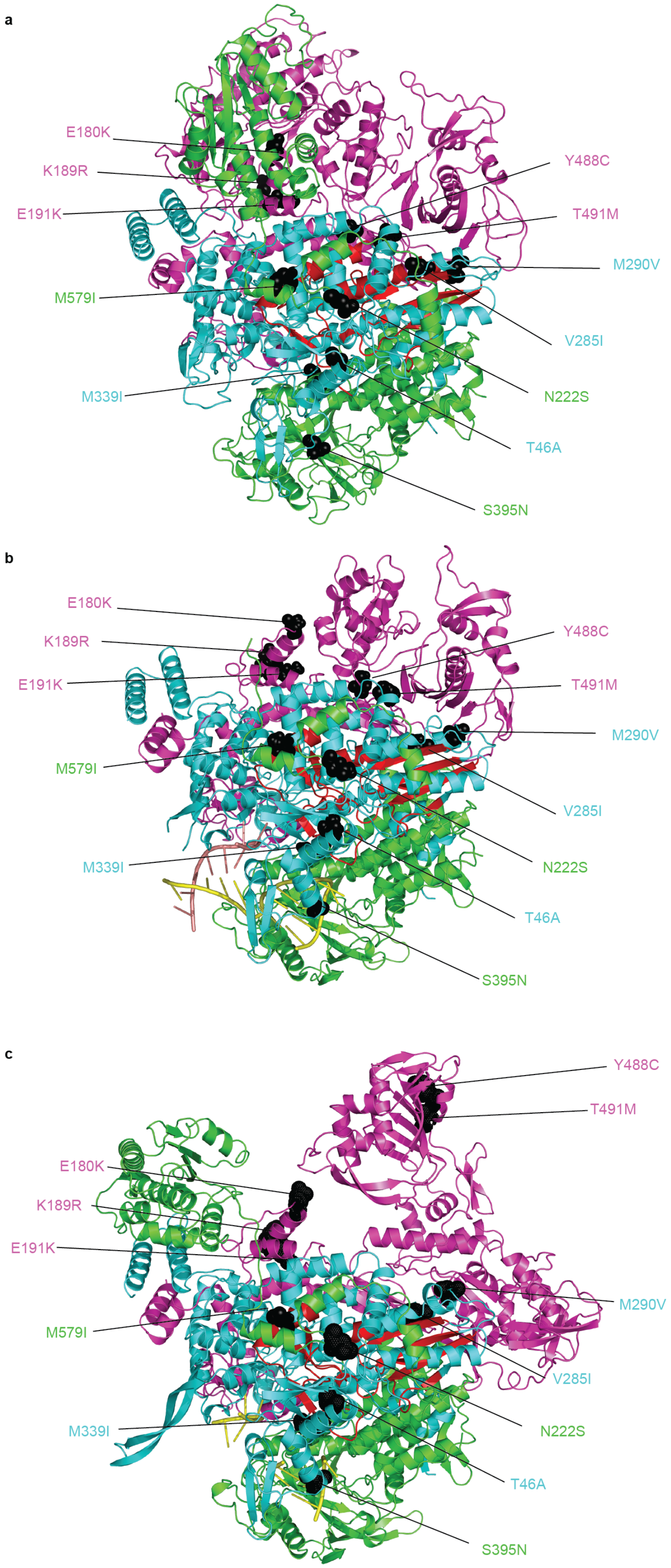
a
E180K
K189R
E191K
M579I
M339I
Y488C
T491M
M290V
V285I
N222S
T46A
S395N
b
E180K
K189R
E191K
M579I
M339I
Y488C
T491M
M290V
V285I
N222S
T46A
S395N
c
Y488C
T491M
E180K
K189R
E191K
M579I
M339I
M290V
V285I
N222S
T46A
S395N

**S5 Figure:** Spatial conservation of 4'-FIU resistance mutations in discrete IAV RdRP structural models. **a-c)** Locations of all 4'-FIU resistance mutations in a CA09 homology model based on the coordinates released for influenza C polymerase (PDBID 5d9a) (a), the 1918 H1N1 influenza A polymerase (PDBID 7ni0) (b), and a bat influenza A polymerase (4wsb) (c). Mutations are shown as black spheres, labels are color-coded by polymerase subunit; PA, green; PB1, cyan; PB2, magenta. The active site for phosphodiester bond formation of the RdRP is shown in red. Homology models were created using SWISS-MODEL, images were created using Pymol.

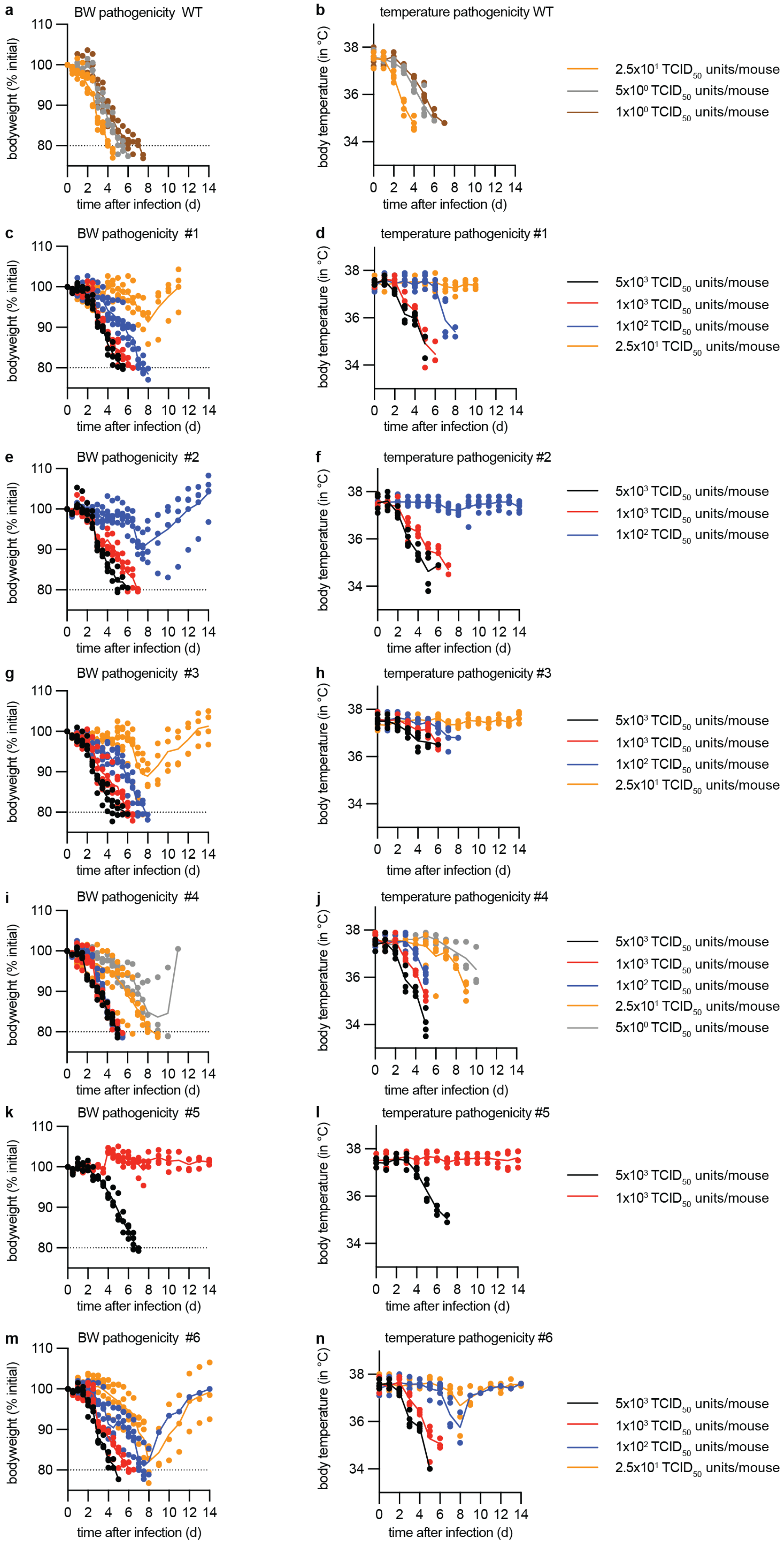
a
BW pathogenicity WT
b
temperature pathogenicity WT
2.5x10¹ TCID50 units/mouse
5x10⁰ TCID50 units/mouse
1x10⁰ TCID50 units/mouse
c
BW pathogenicity #1
d
temperature pathogenicity #1
5x10³ TCID50 units/mouse
1x10³ TCID50 units/mouse
1x10² TCID50 units/mouse
2.5x10¹ TCID50 units/mouse
e
BW pathogenicity #2
f
temperature pathogenicity #2
5x10³ TCID50 units/mouse
1x10³ TCID50 units/mouse
1x10² TCID50 units/mouse
g
BW pathogenicity #3
h
temperature pathogenicity #3
5x10³ TCID50 units/mouse
1x10³ TCID50 units/mouse
1x10² TCID50 units/mouse
2.5x10¹ TCID50 units/mouse
i
BW pathogenicity #4
j
temperature pathogenicity #4
5x10³ TCID50 units/mouse
1x10³ TCID50 units/mouse
1x10² TCID50 units/mouse
2.5x10¹ TCID50 units/mouse
5x10⁰ TCID50 units/mouse
k
BW pathogenicity #5
l
temperature pathogenicity #5
5x10³ TCID50 units/mouse
1x10³ TCID50 units/mouse
m
BW pathogenicity #6
n
temperature pathogenicity #6
5x10³ TCID50 units/mouse
1x10³ TCID50 units/mouse
1x10² TCID50 units/mouse
2.5x10¹ TCID50 units/mouse
bodyweight (% initial)
body temperature (in °C)
time after infection (d)

**S6 Figure:** Clinical signs in mice enrolled in pathogenesis assessment. **a, c, e, g, I, k, m)** Bodyweight normalized to weight at the time of infection for parental recCA09 and the rebuilt resistance lineages #1-6. Dashed line, predefined humane endpoint. **b, d, f, h, j, l, n)** Rectal temperature measured once daily. Symbols represent individual animals, lines connect data means; n=4-5.

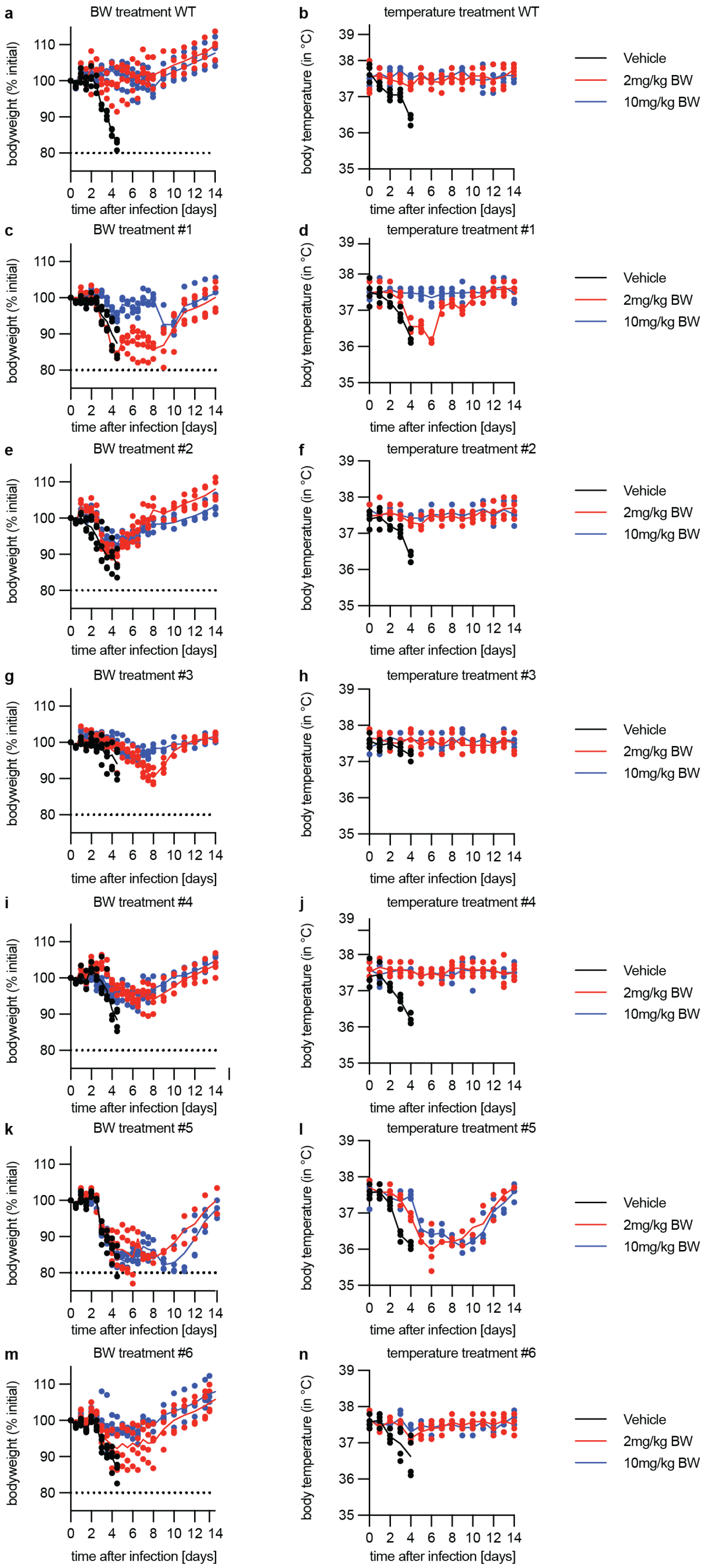
a
BW treatment WT
b
temperature treatment WT
c
BW treatment #1
d
temperature treatment #1
e
BW treatment #2
f
temperature treatment #2
g
BW treatment #3
h
temperature treatment #3
i
BW treatment #4
j
temperature treatment #4
k
BW treatment #5
l
temperature treatment #5
m
BW treatment #6
n
temperature treatment #6
bodyweight (% initial)
body temperature (in °C)
time after infection [days]
Vehicle
2mg/kg BW
10mg/kg BW

**S7 Figure:** Clinical signs in mice enrolled in efficacy testing. **a, c, e, g, I, k, m)** Bodyweight normalized to weight at the time of infection for parental recCA09 and the rebuilt resistance lineages #1-6. Dashed line, predefined humane endpoint. **b, d, f, h, j, l, n)** Rectal temperature measured once daily. Symbols represent individual animals, lines connect data means; n=4.

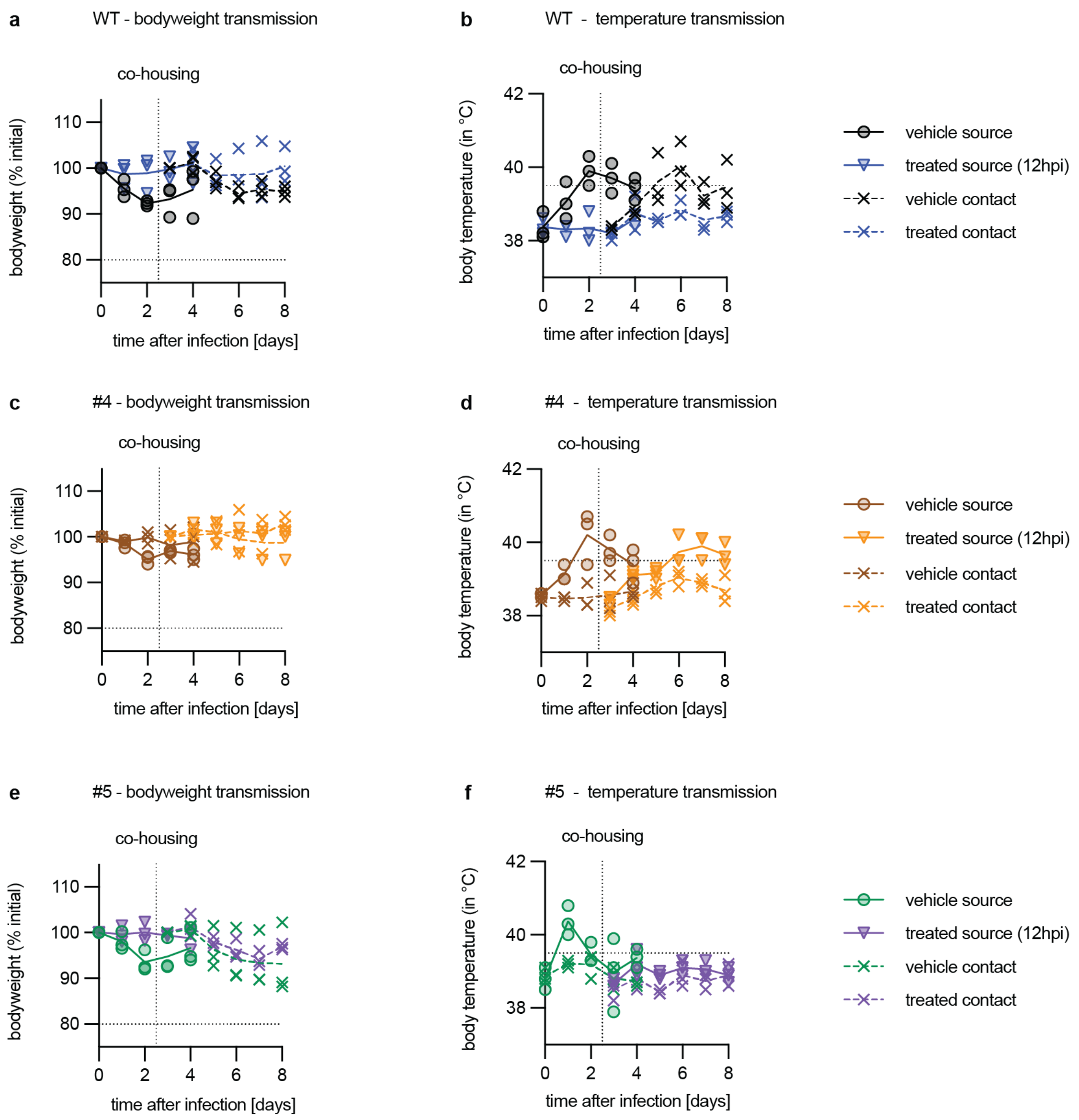


**S8 Figure:** Clinical signs in mice enrolled in transmission studies. **a, c, e:** Bodyweight normalized to weight at the time of infection for parental recCA09 (a) and resistance lineages #4 (c) and #5 (e). Dashed line, predefined humane endpoint. **b, d, f)** Rectal temperature of recCA09 (b) and resistance lineages #4 (d) and #5 (f) measured once daily. Dashed line, onset of fever (39.5□C). Symbols represent individual animals, lines connect data means; n=3.

**S1 Data:** RAVA plots covering virus adaptations. Data can be found at https://github.com/greninger-lab/Lieber_IAV_4FIU_Supplemental_Data_1-4/tree/main/S1_Data/adaptation_rava_plots

**S2 Data:** RAVA plots covering *in vitro* virus fitness tests. Data can be found at https://github.com/greninger-lab/Lieber_IAV_4FIU_Supplemental_Data_1-4/tree/main/S2_Data/fitness_rava_plots

**S3 Data:** RAVA plots covering virus populations isolated from infected mice. Data can be found at https://github.com/greninger-lab/Lieber_IAV_4FIU_Supplemental_Data_1-4/tree/main/S3_Data/mouse_rava_plots

**S4 Data:** RAVA plots covering virus populations isolated from infected ferrets. Data can be found at https://github.com/greninger-lab/Lieber_IAV_4FIU_Supplemental_Data_1-4/tree/main/S4_Data/ferret_rava_plots

**S5 Data:** All quantitative raw data

**S6 Data:** All statistical analyses
